# Whole transcriptome data of uninfected and *Nosema ceranae*-infected midguts of eastern honeybee workers

**DOI:** 10.1101/2020.03.13.990697

**Authors:** Huazhi Chen, Dingding Zhou, Yu Du, Cuiling Xiong, Yanzhen Zheng, Dafu Chen, Rui Guo

## Abstract

*Apis cerana cerana* is a subspecies of eastern honeybee, *Apis cerana. Nosema ceranae* is a widespread fungal parasite of honeybee, causing heavy losses for beekeeping industry all over the world. In this article, total RNA of normal midguts (AcCK1, AcCK2) and *N. ceranae*-infected midguts of *A. c. cerana* workers at 7 d and 10 d post inoculation (AcT1, AcT2) were respectively isolated followed by strand-specific cDNA library construction and next-generation RNA sequencing. In tolal, 56270223688, 44860946964, 78991623806, and 92712308296 raw reads were derived from AcCK1, AcCK2, AcT1 and AcT2, respectively. Following strict quality control, 54495191388, 43570608753, 76708161525, and 89467858351 clean reads were obtained, with Q30 value of 95.80%, 95.99%, 96.07% and 96.04%, and GC content of 44.20%, 43.44%, 44.83% and 43.63%, respectively. The raw data were submitted to the NCBI Sequence Read Archive database and connected to BioProject PRJNA562784. These data offers a valuable resource for deep investigation of mechanisms underlying eastern honeybee responding to *N. ceranae* infection and host-fungal parasite interaction during microsporidiosis.

**Value of the Data:**

- Current dataset offers a valuable resource for exploring mRNAs, lncRNAs and circRNAs involved in response of *A. c. cerana* worker to *N. ceranae* infection.
- The accessible data can be used to investigate differential expression pattern and regulatory network of non-coding RNAs in *A. c. cerana* workers’ midguts responding to *N. ceranae* challenge.
- This data will enable a better understanding of the molecular mechanism regulating eastern honeybee-*N. ceranae* interaction.

## Data

The shared datasets were derived from strand-specific cDNA library-based RNA-seq of un-infected (AcCK1 and AcCK2) and *N. ceranae*-infected (AcT1 and AcT2) *A. c. cerana* workers’ midguts [1]. Totally, 56270223688, 44860946964, 78991623806, and 92712308296 raw reads were gained from AcCK1, AcCK2, AcT1 and AcT2, respectively (Table 1). In addition, 54495191388, 43570608753, 76708161525, and 89467858351 clean reads were obtained after strict quality control, with Q30 value of 95.80%, 95.99%, 96.07%, and 96.04%, and GC content of 44.20%, 43.44%, 44.83%, and 43.63%, respectively (Table 1). Pearson correlation coefficients among three replicas in each group were above 0.8772 (Fig. 1). The raw data were submitted to the NCBI Sequence Read Archive (SRA) database and connected to BioProject PRJNA562784.

**Table 1.**
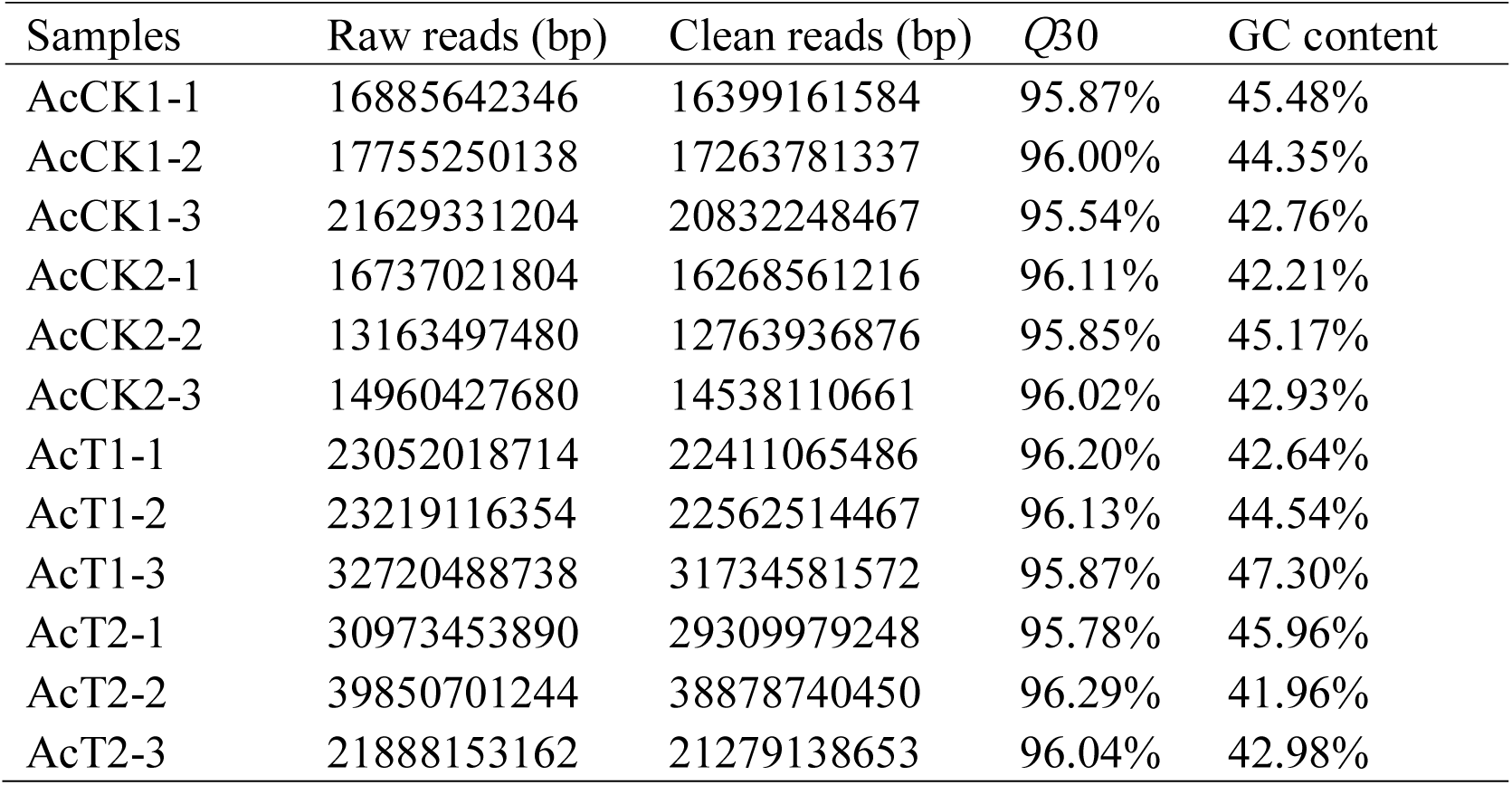
Overview of strand-specific cDNA library-based RNA-seq data.

**Fig. 1.**
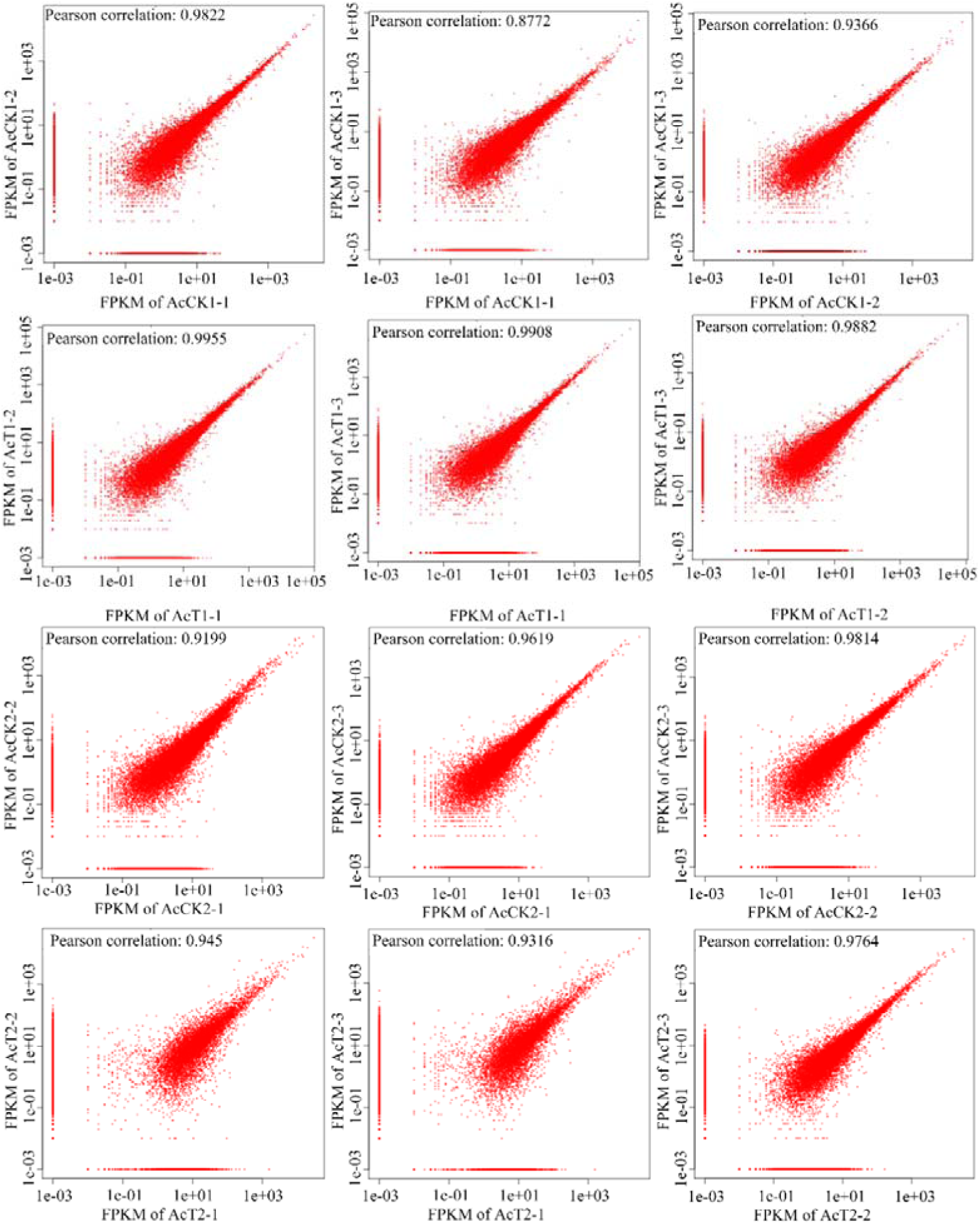
Pearson correlation coefficients among different replicas within every un-infected and *N. ceranae*-infected groups.

## Experimental Design, Materials, and Methods

### 2.1 N. ceranae spore purification

*N. ceranae* spores were previously purified from midguts of *Apis mellefera ligustica* workers infected by *N. ceranae* [2]. The spore suspension was freshly prepared before use.

### 2.2 Experimental design and sample collection

Frames of sealed brood obtained from a healthy colony of *A. c. cerana* located in the teaching apiary of College of Animal Sciences (College of Bee Science), Fujian Agriculture and Forestry University were reared in an incubator at 34 ± 0.5 °C, 50% RH to provide newly emerged *Nosema*-free honeybees. The emergent workers were carefully removed and confined to cages in groups (n=20), and kept in the incubator at 32 ± 0.5 °C, 50% RH.

The workers were fed *ad libitum* with a f sucrose solution (50% w/v in water), and one day after eclosion, workers in treatment groups were starved for 2 h and 20 workers per group were each immobilized and then fed with 5 uL of 50% sucrose solution containing 1×10^6^ fungal spores. Those workers that did not consume the total amount of sucrose solution were discarded. After inoculation, workers were isolated for 30 min in vials in the growth chamber to ensure that the sucrose solution was not transferred among individuals and the entire dosage was ingested. Workers in control groups were inoculated in an identical manner using a 50% sucrose solution (w/w in water) without *N. ceranae* spores. Three replicate cages of 20 honeybees each were used in treatment and control groups. Each cage was checked every 24 h and any dead bees removed. *N. ceranae*-infected and un-infected workers’ midguts were respectively harvested at 7 d or 10 d post inoculation (dpi), immediately frozen in liquid nitrogen and kept at −80 °C until RNA sequencing. *N. ceranae*-infected groups at 7 dpi and 10 dpi with sucrose solution containing *N. ceranae* spores were termed as AcT1 (AcT1-1, AcT1-2, AcT1-3) and AcT2 (AcT2-1, AcT2-2, AcT2-3); un-infected groups at 7 dpi and 10 dpi with sucrose solution without *N. ceranae* spores were termed as AcCK1 (AcCK1-1, AcCK1-2, AcCK1-3) and AcCK2 (AcCK2-1, AcCK2-2, AcCK2-3).

### 2.3. Strand-specific cDNA library construction and illumina sequencing

Firstly, total RNA of the six midgut samples from *N. ceranae*-infected groups and six midgut samples from un-infected groups were respectively extracted using Trizol (Life Technologies) following the manufacturer’s protocol, and examined via 1% agarose gel eletrophoresis. Secondly, rRNAs were removed to retain mRNAs and non-coding RNAs (ncRNAs), which were fragmented into short fragments with fragmentation buffer (Illumina, USA) followed by reverse transcription into cDNA with random primers. Thirdly, second-strand cDNA were synthesized by dNTP (dUTP instead of dTTP), DNA polymerase I, RNase H, and buffer. Fourthly, the cDNA fragments were purified using QiaQuick PCR extraction kit (QIAGEN, Germany), end repaired, poly(A) added, and ligated to Illumina sequencing adapters, followed by digestion of the second-strand cDNA with UNG (Uracil-N-Glycosylase) (Illumina, USA). Finally, the digested products were size selected by agarose gel electrophoresis, PCR amplified, and sequenced on Illumina HiSeq™ 4000 platform (Illumina, USA) by Gene Denovo Biotechnology Co. (China). The raw sequencing data produced in this article are available in NCBI SRA database and connected to BioProject PRJNA562784.

### 2.4. Quality control of raw reads and mapping of clean reads

Firstly, raw reads were filtered by removing reads containing adapters, more than 10% of unknown nucleotides (N), and more than 50% of low quality bases to obtain high quality clean reads. Quality control of transcriptome data is showed in Table 1. Secondly, the filtered raw reads were mapped to ribosome RNA (rRNA) database using short reads alignment tool Bowtie2 [3]. Thirdly, the mapped reads were removed and the remaining reads were used in assembly and further analysis. The rRNA-removed reads of each sample were then mapped to *Apis cerana* genome (assembly ACSNU-2.0) using TopHat2 (version 2.0.3.12) [4] following alignment parameters: (1) maximum read mismatch is two; (2) the distance between mate-pair reads is 50 bp; (3) the error of distance between mate-pair reads is ±80 bp.

### 2.5. Transcript assembly

Transcripts were assembled using Cufflinks [5]. The program reference annotation based transcripts (RABT) was preferred. Cufflinks constructed faux reads based on reference to make up for the influence of low coverage sequencing. During the last step of assembly, all of the reassembles fragments were aligned with reference genes followed by removel of similar fragments. Cuffmerge was used to merge transcripts from different replicas of a group into a comprehensive set of transcripts, and the transcripts from multiple groups were then merged into a final set of transcripts. Pearson correlation coefficients between every biological replicas in each sample group were calculated and presented in Fig. 1.

## Acknowledgments

This work was supported by the Earmarked Fund for Modern Agro-industry Technology Research System (CARS-44-KXJ7), the Science and Technology Planning Project of Fujian Province (2018J05042), the Education and Scientific Research Program Fujian Ministry of Education for Young Teachers (JAT170158), the Outstanding Young Scientific Research Talents Program of Fujian Agriculture and Forestry university (xjq201814), and the Fujian Agriculture and Forestry University Science and Technology Innovation Special Fund (CXZX2017342, CXZX2017343).

